# *De novo* assembly and annotation of the larval transcriptome of two spadefoot toads widely divergent in developmental rate

**DOI:** 10.1101/421446

**Authors:** H. Christoph Liedtke, Jèssica Gómez Garrido, Marta Gut, Anna Esteve-Codina, Tyler Alioto, Ivan Gomez-Mestre

## Introduction

Most amphibian species exhibit a complex life-cycle including two or more life stages separated by an ontogenetic switch point such as hatching or metamorphosis. Adaptations to divergent environments can require the modification of the timing of such switch points and the relative investment in growth and differentiation between subsequent stages [1]. Such alterations of developmental trajectories, however, often have substantial repercussions at several organismal levels, from physiology to morphology and even genomic structure. Adaptive divergence in developmental rate tracking aquatic habitats of different duration in spadefoot toads is a well-known example of this. Spadefoot toads from Europe and northern Africa typically have tadpoles that grow to be quite large over a long larval period, but can otherwise accelerate development and precipitate metamorphosis if at risk of pond drying, whereas north American spadefoot toads tend to have smaller tadpoles that develop faster and are less capable of further developmental acceleration [2]. At each end of this spectrum we find *Pelobates cultripes*, distributed throughout most of the Iberian Peninsula and southern France, and *Scaphiopus couchii*, distributed across southwestern USA to northern Mexico. *Pelobates cultripes* larvae grow quite large (up to > 16 g) and can take up to 6 months to reach metamorphosis, whereas *S. couchii*’s tadpoles are much smaller (1.5-2 g) and can develop to metamorphosis in as little as 8 days. Such developmental acceleration is rather energetically demanding and requires a substantial increase in metabolic activity [3], hence incurring in oxidative stress [4]. Precipitating metamorphosis alters growth and developmental trajectories non-isometrically for different parts of the body, causing metamorphs to not only be smaller but also to have relatively shorter limbs [5-7]. Developmental acceleration is achieved through neuroendocrine regulation mainly resulting in increased corticosterone and thyroid hormone levels [3, 8], as well as through differential expression of hormone receptors [4]. Interestingly, the canalized fast development of *S. couchii* to a large extent mirrors the environmentally-induced accelerated state of the more plastic *P. cultripes* [3].

At the genomic level, evolutionary divergences in developmental rate seem to leave a big imprint on whole genomes with some studies showing that fast developmental rates are often associated with smaller genome sizes [9, 10]. The rule also holds true for amphibians, whether at a large macroevolutionary scale (Liedtke et al. *in press*) or focused on specific species groups [11]. Spadefoot toads present broad differences in developmental rate across species, which are consequently also reflected in large differences in genome size [12]: slow developing *Pelobates cultripes* has a large genome (∼3.9 Gbp), whereas fast developing *S. couchii* has only about one third its size (∼1.5 Gbp). Here we present a first description of the transcriptomes of these species at the onset of metamorphosis to explore the potential consequences of such dramatic divergence in their genomes and to uncover the transcriptomic basis of their differences in developmental rate.

The NCBI Transcriptome Shotgun assemblies database currently lists transcriptome assemblies for 26 species of amphibians and of those, only four are larval phase transcriptomes: *Rhinella marina* [13], *Microhyla fissipes* [14], *Lithobates catesbeiana* and *Xenopus laevis* [15]. The addition of transcriptomes for the larval phases of two more species, especially as they represent a distinct evolutionary lineage, is therefore a significant contribution to the current knowledgebase.

## Methods

### Sample collection, total RNA extraction and sequencing

Three egg clutches of *P. cultripes* were collected from a natural pond in Doñana National Park, southwestern Spain, brought to a walk-in chamber in the laboratories of Doñana Biological Station (EBD-CSIC) and placed in a plastic tray with carbon-filtered dechlorinated tap water with aerators to ensure adequate oxygenation. Another three clutches of *Scaphiopus couchii* were obtained from adult pairs kept in the laboratory at EBD-CSIC. Adults were hormonally stimulated to breed by intraperitoneally injecting 20–100 µL of 1 µg/100 µL GnRH agonist (des-Gly, [D-His(Bzl)]- luteinizing hormone releasing hormone ethylamide, Sigma). Upon hatching, we transferred tadpoles from each clutch of each species to 3 L plastic containers with dechlorinated tap water where they were individually kept under standard conditions of 24 °C, 12:12 L:D photoperiod, *ad libitum* food supply consisting of finely powdered rabbit chow. As tadpoles reached Gosner stage 35 in their development [16], we euthanized twelve individuals per species via MS-222 overdose, eviscerated them to avoid interreferences from faecal material, and snap-froze them in liquid nitrogen. We extracted whole-body total RNA from each tadpoles using Trizol reagent following the manufacturer’s protocol (Invitrogen). Total RNA was assayed for quantity and quality using Qubit® RNA HS Assay (Life Technologies) and RNA 6000 Nano Assay on a Bioanalyzer 2100.

The RNASeq libraries were prepared from total RNA using the TruSeq®Stranded mRNA LT Sample Prep Kit (Illumina Inc., Rev.E, October 2013). Briefly, 500ng of total RNA was used as the input material and was enriched for the mRNA fraction using oligo-dT magnetic beads. The mRNA was fragmented in the presence of divalent metal cations. The second strand cDNA synthesis was performed in the presence of dUTP instead of dTTP, this allowed to achieve the strand specificity. The blunt-ended double stranded cDNA was 3′adenylated and Illumina indexed adapters were ligated. The ligation product was enriched with 15 PCR cycles and the final library was validated on an Agilent 2100 Bioanalyzer with the DNA 7500 assay.

Each library was sequenced using TruSeq SBS Kit v3-HS, in paired end mode with the read length 2×76bp. We generated on average 38 million paired-end reads for each sample in a fraction of a sequencing lane on HiSeq2000 (Illumina) following the manufacturer’s protocol. Images analysis, base calling and quality scoring of the run were processed using the manufacturer’s software Real Time Analysis (RTA 1.13.48) and followed by generation of FASTQ sequence files by CASAVA 1.8.

### *Assembling* de novo *transcriptomes of* P. cultripes *and* S. couchii

Quality of raw reads was inspected using FASTQC (https://www.bioinformatics.babraham.ac.uk/projects/fastqc/) and MULTIQC [17]. Assembly was performed using Trinity v2.4.0 [18] for the two species separately. Reads from all samples per species were combined, trimmed (using default Trimmomatic settings SLIDINGWINDOW:4:5 LEADING:5 TRAILING:5 MINLEN:25)[19] and normalized using *in silico* normalization with default Trinity settings (flags used: --trimmomatic --normalize_max_read_cov 50).

### Assessment of transcriptome quality and completeness

Transcriptome quality in terms of read representation was evaluated by mapping the normalized reads (pairs only) back onto the transcriptome using Bowtie2 v2.3.2 [20]. Completeness in terms of gene content was assessed using BUSCO v3.0.2 [21] with the tetrapoda-odb9 database as a reference as well as by running blastx (E-value cut off E≤1e^-20^) against both the SwissProt database (downloaded on 01.11.2017) and the *Xenopus tropicalis* proteome (Ensemble JGI 4.2; downloaded on 03.11.2017) with a stringent Evalue criteria of ≤ 1e^-20^. The count of full-length transcripts with blastx hits was based on grouped high scoring segment pairs per transcript to avoid multiple fragments per transcript aligning to a single protein sequence.

### Functional annotation

We used Trinotate v3.0 (https://trinotate.github.io/) to annotate the transcriptome. This involves finding similarities to known proteins by querying transcripts against the Swissprot database (accessed in June 2018) [22] (blastx with a cut-off of E≤1e^-5^). Moreover, likely coding regions were detected with TransDecoder (https://github.com/TransDecoder) and resulting protein products (coding sequence; CDS) were matched against both the complete Swissprot database and a subset including only vertebrate genes, using blastp (Evalue≤1e^-5^), and a conserved protein domain search was conducted using Hmmr (http://hmmer.org/) on the Pfam database [23]. SignalP v4.1 [24] and TmHMM v2.0 (http://www.cbs.dtu.dk/services/TMHMM/) were used to predict signal peptides and transmembrane regions respectively. Finally, gene ontology identifiers were assigned to transcripts based on available annotations from best-matching Swissprot entries. Trinotate also provides KEGG (Kyoto Encycopedia of Genes and Genomes; http://www.genome.jp/kegg/) and EggNOG [25] annotations. Exploring the Trinotate output was facilitated using the TrinotateR R package (https://github.com/cstubben/trinotateR).

The PANTHER classification scheme [26] for *Xenopus tropicalis* was used to organize gene function and ontology using the *Xenopus* Ensembl protein identifiers recovered for the quality assessment step above. Using the PANTHER web server, we performed both functional classifications and a statistical overrepresentation test (with default settings), to investigate which genes are significantly (p<0.05) over or under represented in our transcriptomes compared to the *X. tropicalis* reference.

### Orthologous genes

Orthofinder v2.2.3 [27] was used to find orthogolous genes across the two species. Orthofinder was run with default settings, taking the transdecoder predicted CDS of both *P. cultripes* and *S. couchii* as the input, as well as the proteome of *X. tropicalis* (JGI 4.2) to provide context (as an ‘outgroup’).

## Results and Discussion

### Transcriptome comparison and quality assessment

The twelve *P. cultripes* samples consisted of 30.5-43.3 million, 101bp paired-end reads (888.3 million reads in total) pooling to 84.2 million post-normalization pair-end reads used for the assembly (10.5% of total). Trinity generated 753,223 transcript contigs with median length 362, of which 428 406 clustered into ‘genes’ (transcript clusters with shared sequence content; Table 1). Bowtie2 mapped 83.96% of the reads back onto the transcriptome (Supporting Data 1). In comparison, the *S. couchii* samples consisted of 32.1-53.9 million, 101bp reads (958.8 million reads in total) with 84.4 million post-normalization pair-end reads used in for the final assembly (9.19%). 657,280 transcripts were generated by Trinity with a median length of 432bp clustering into 381,135 ‘genes’ (Table 1). Bowtie2 mapped 90.71% of the reads back onto the transcriptome.

**Table 1:**
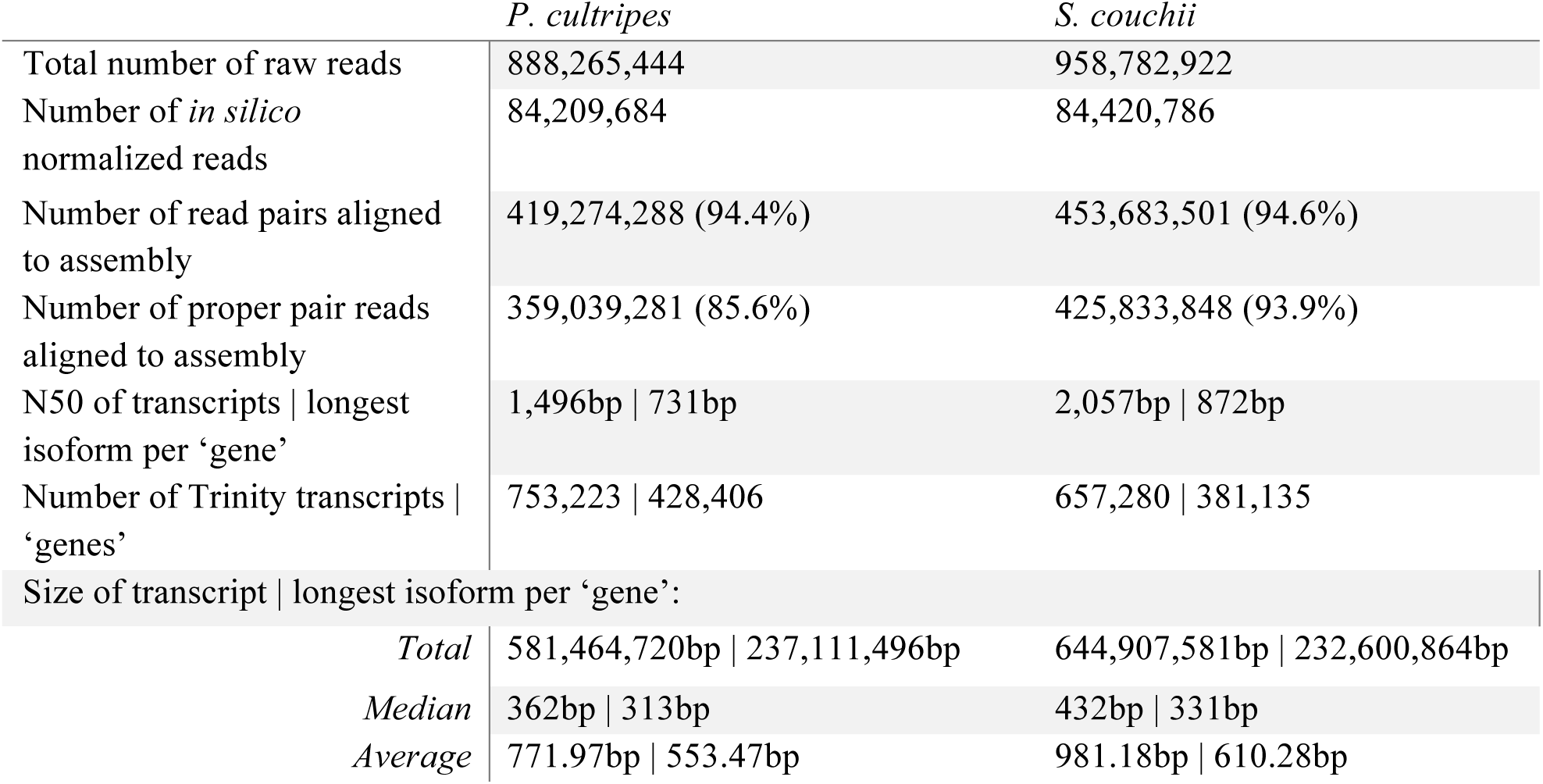
Transcriptome assembly statistics for both tadpole species. Summaries for Trinity outputs are given both at the transcript and at the ‘gene’ level.

The BUSCO results support near-complete gene sequence information for 89.7% of genes in the *P. cultripes* transcriptome with only 7.4% of the genes being fragmented and 2.9% missing. The quality of the *S. couchii* assembly was similar with 86.6% complete sequence information, 10.5% fragmented genes and 2.9% missing (Supporting Data 2).

Querying the Trinity assembly against both the Swissprot database and the *X. tropicalis* proteome (using blastx) revealed large numbers of fully reconstructed coding transcripts, with 13,645 Swissprot proteins and 12,715 *X. tropicalis* proteins represented by nearly full-length transcripts (>80% alignment coverage) in the *P. cultripes* assembly, and 14,429 Swissprot proteins and 12,216 *X. tropicalis* proteins in the *S. couchii* assembly (Figure 1; Supporting Data 3).

**Figure 1:**
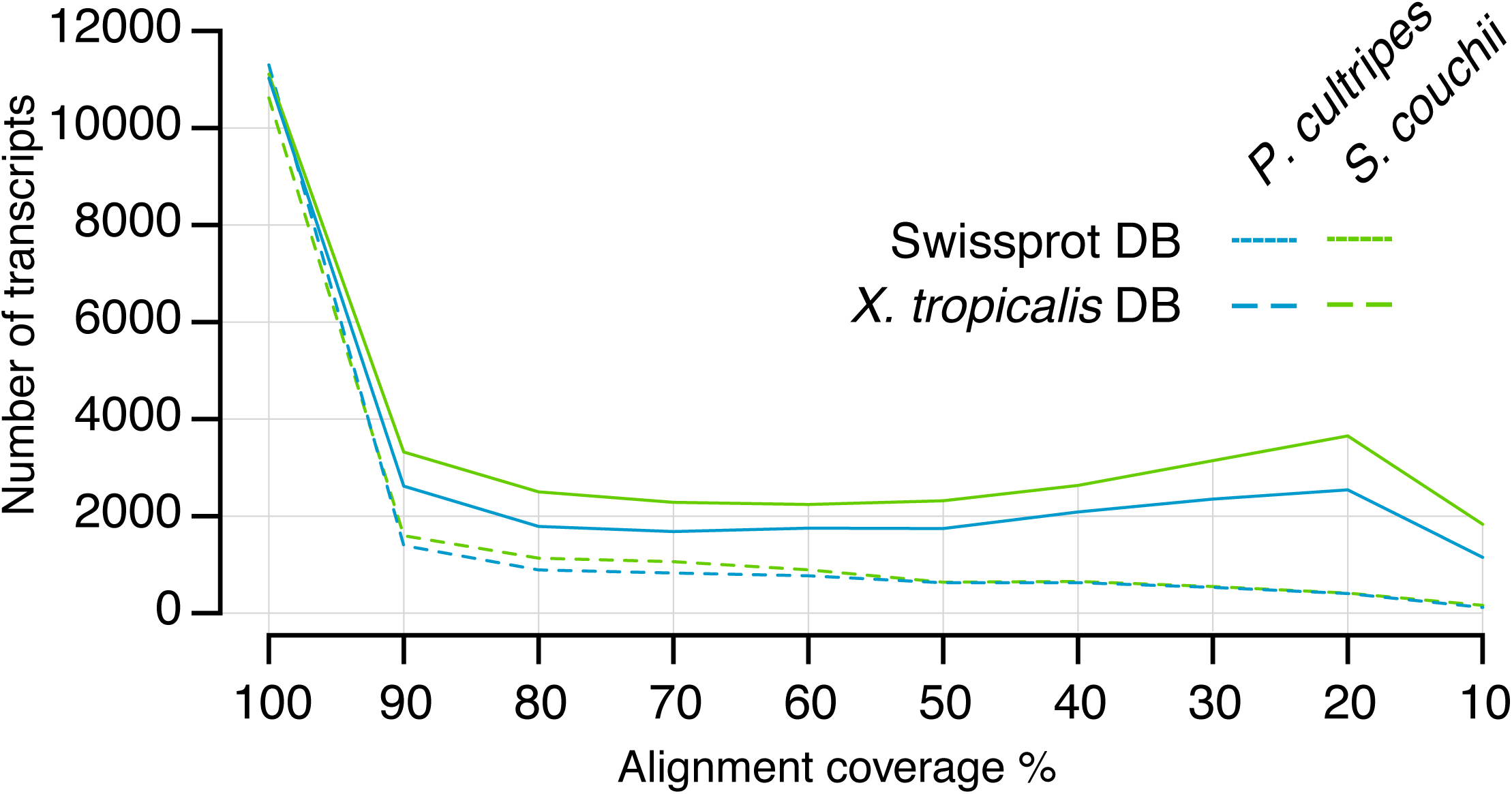
Number of transcripts (using grouped highest scoring segment pairs) per alignment coverage bins when querying against the SwissProt and *Xenpus tropicalis* proteome sequence databases.

### Functional annotation

Gene annotation via the Trinotate pipeline is useful for providing biological context to the assembled transcriptomes (Trinotate tables available as Supporting Data 4). Querying (using blastx) the SwissProt database with the trinity assembly allowed for the annotation of 162,031 *P. cultripes* and 204,646 *S. couchii* transcripts. Gene Ontology (GO) derived from these hits resulted in 18,585 unique (out of a total of 1,626,015) GO annotations for *P. cultripes* and 19,917 unique (out of a total 2,155,849) GO annotations for *S. couchii*. The most abundant GO terms per ontology (based on number of corresponding Trinity ‘genes’) for both species were largely comparable for cellular components (CC) and molecular function (MF) ontologies, but different for biological processes (BP), with genes related to DNA recombination, RNA mediated transposition and DNA integration being abundant in *P. cultripes* and notably less so (not in the top ten most abundant genes) in *S. couchii* (Figure 2).

**Figure 2.**
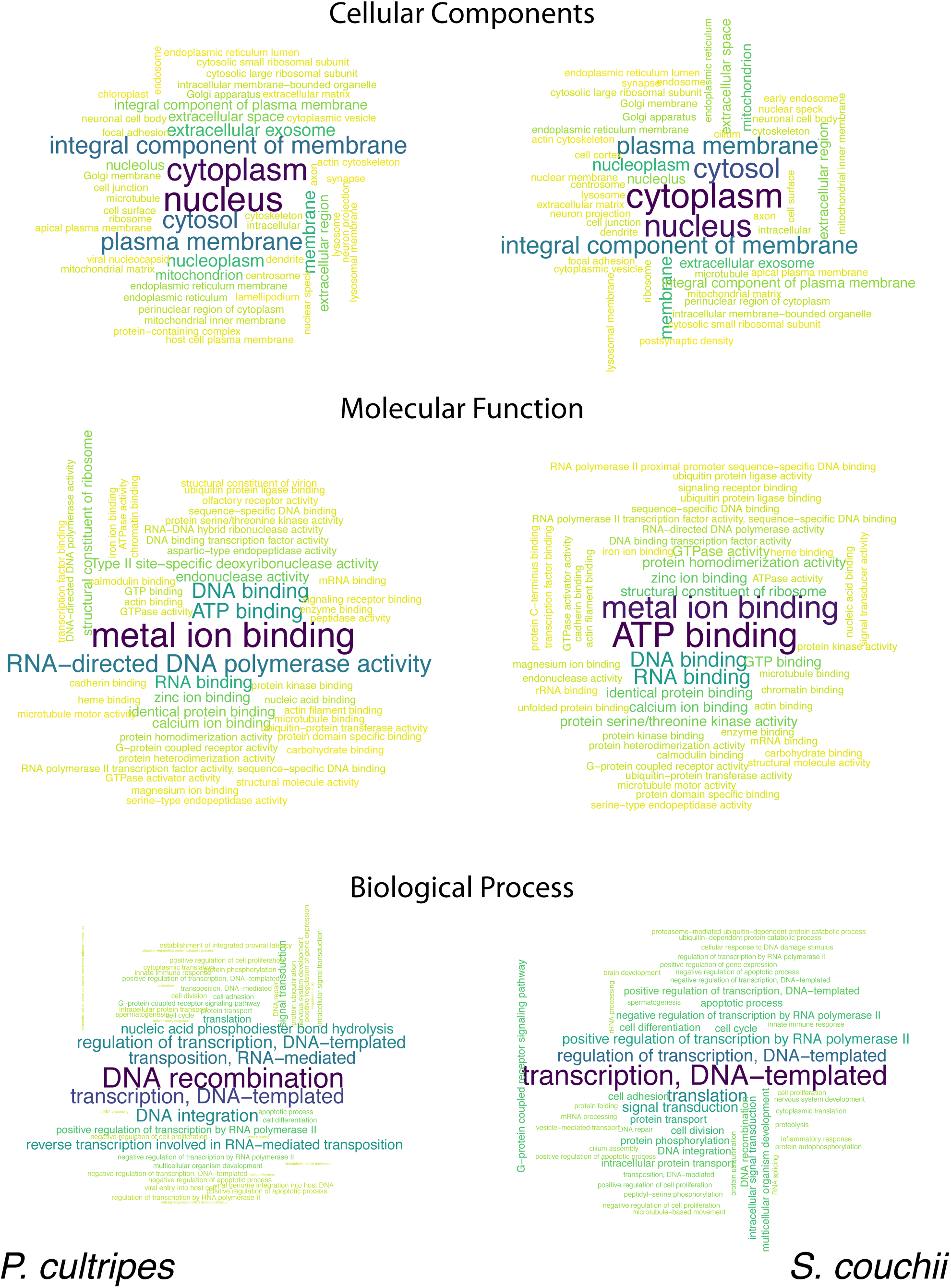
Wordclouds of the 50 most abundant GO terms per ontology per species. Size and colour (large to small, dark to light) is relative to the number of Trinity ‘genes’ that are associated with each GO term.

The PANTHER GO-slim classification system designed for *Xenopus tropicalis* provides a curated, functional classification scheme of GO terms and allows for relative over or underrepresentation of terms to be assessed in relation to the reference (in this case *X. tropicalis*) database. For 17,488 unique *X. tropicalis* genes recovered for *P. cultripes* and 17,717 for *S. couchii*, 13,658 and 13,622 could be mapped to PANTHER genes respectively. The majority of PANTHER terms are overrepresented in both species in comparison to the *X. tropicalis* reference, including the most extensively represented GO terms for each of the three ontologies (Figure 3). These are comparable across the two species (Figure 3) with most of the transcriptomes being related to cell parts and organelles (cellular components; CC), binding and catalytic activity (molecular function; MF) and cellular and metabolic processes (biological processes; BP). Genes related to receptor and transporter activity (MF) are underrepresented in both transcriptomes, as are genes relevant for biological regulation (BP) in *P. cultripes* and response to stimulus (BP) in *S. couchii*. Barcharts showing over and underrepresentations of each PANTHER term per species per onotolgy are provided as supporting data (Supporting Data 5).

**Figure 3.**
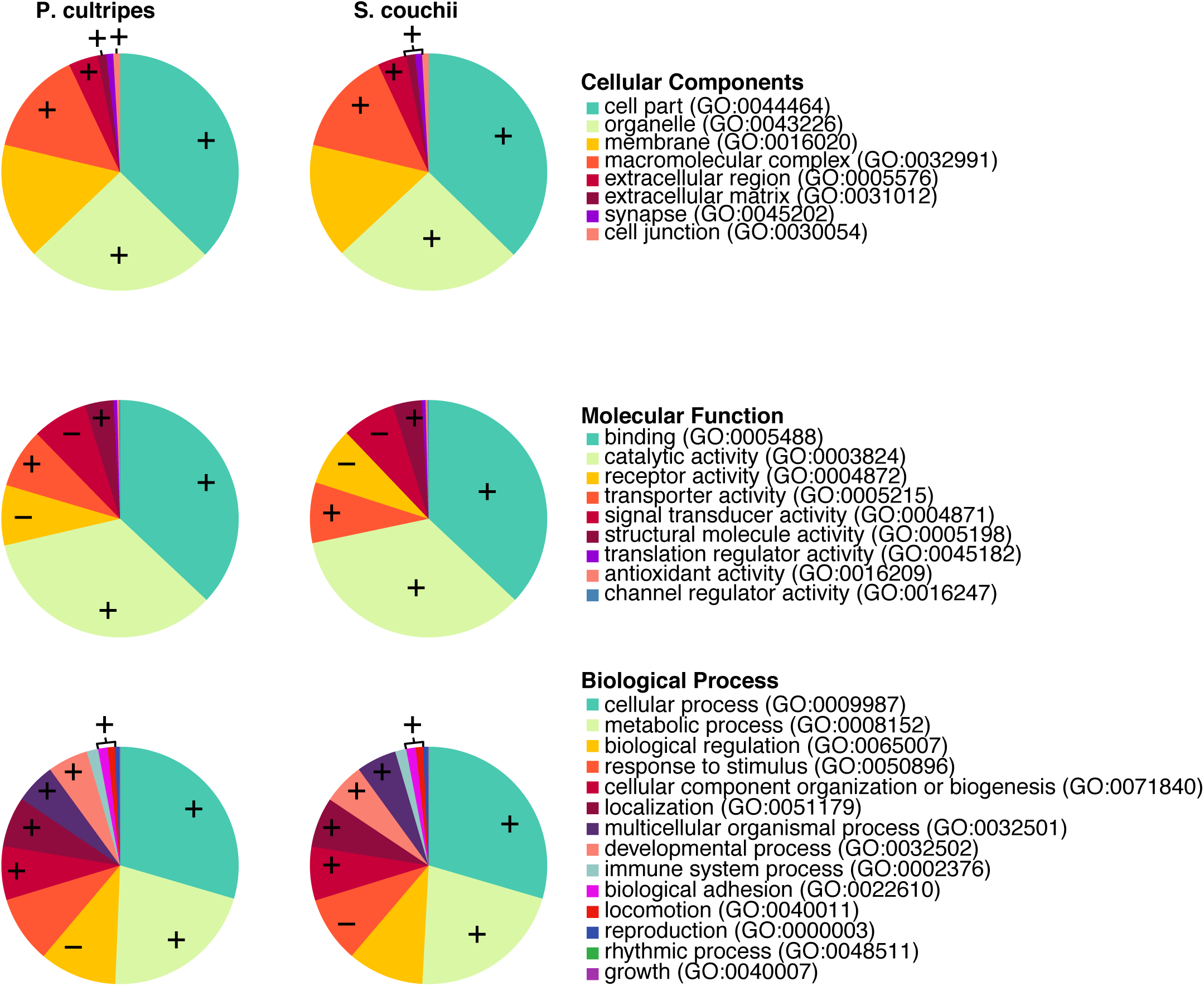
PANTHER functional classification of transcriptomes. Wedge size reflect number of unique genes per category and +/- annotations specify significant over/under representation of the GO-slim term compared to the *X. tropicalis* reference database.

TransDecoder recovered fewer candidate coding regions for *P. cultripes* (154,906) than for *S. couchii* (175,331; Table 2). This corresponds to 36.2% and 46.1% of the Trinity-identified ‘genes’ for *P. cultripes* and *S. couchii* respectively. Homology searches using blastp against the entire Swissprot database was able to annotate 108,881 and 132,578 of these, and 107,199 and 130,847 when searching the vertebrates-only database (Table 2). Of the sequences with vertebrate gene hits, 24,327 and 27,309 unique vertebrate swissprot proteins were identified for *P. cultripes* and *S. couchii* (genes with unique UniProtKB-IDs). Of these, the two species share 56.5% (18,651 proteins), with 17.2% being unique to *P. cultripes* (5,676 proteins) and 26.2% unique to *S. couchii* (8,658 proteins). Similarly, the number of hits of candidate coding regions against other databases including pfam, signalP, tmHMM, KEGG and EggNOG was greater for *S. couchii* than *P. cultripes* (Table 2).

**Table 2:**
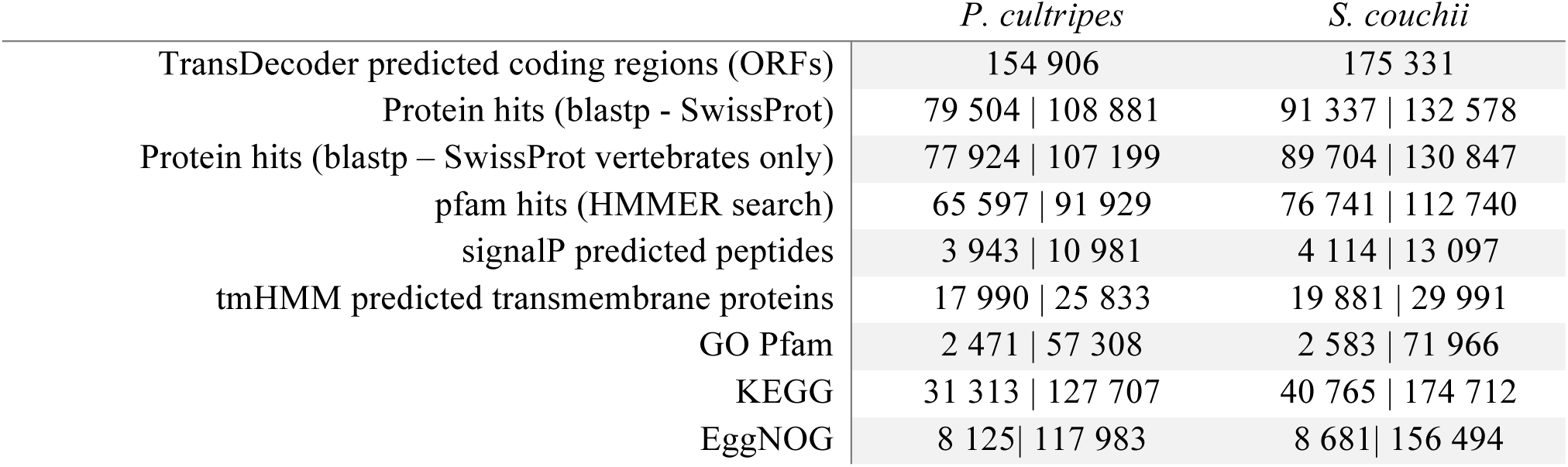
Number of unique | total TransDecoder-predicted candidate genes with annotations via different search tools and databases (summary of Trinotate results).

### Orthologous genes

OrthoFinder assigned 183,893 transdecoder-predicted CDS (52.1% of total, from hereon ‘genes’) to 27,111 orthogroups. Almost all of the *X. tropicalis* genes could be assigned to orthogroups (96.5%), compared to 53.6% of *P. cultripes* genes and 45.7% of *S. couchii* genes (Figure 4a). This could suggest that our transcriptomes represent large numbers of interesting genes not yet represented in the *X. tropicalis* transcriptome, but it is important to note that OrthoFinder may be sensitive to the large number of fragments in *de novo* transcriptome assemblies (compared to its designed use for genome assemblies) and to the number of species included in the analysis.

**Figure 4.**
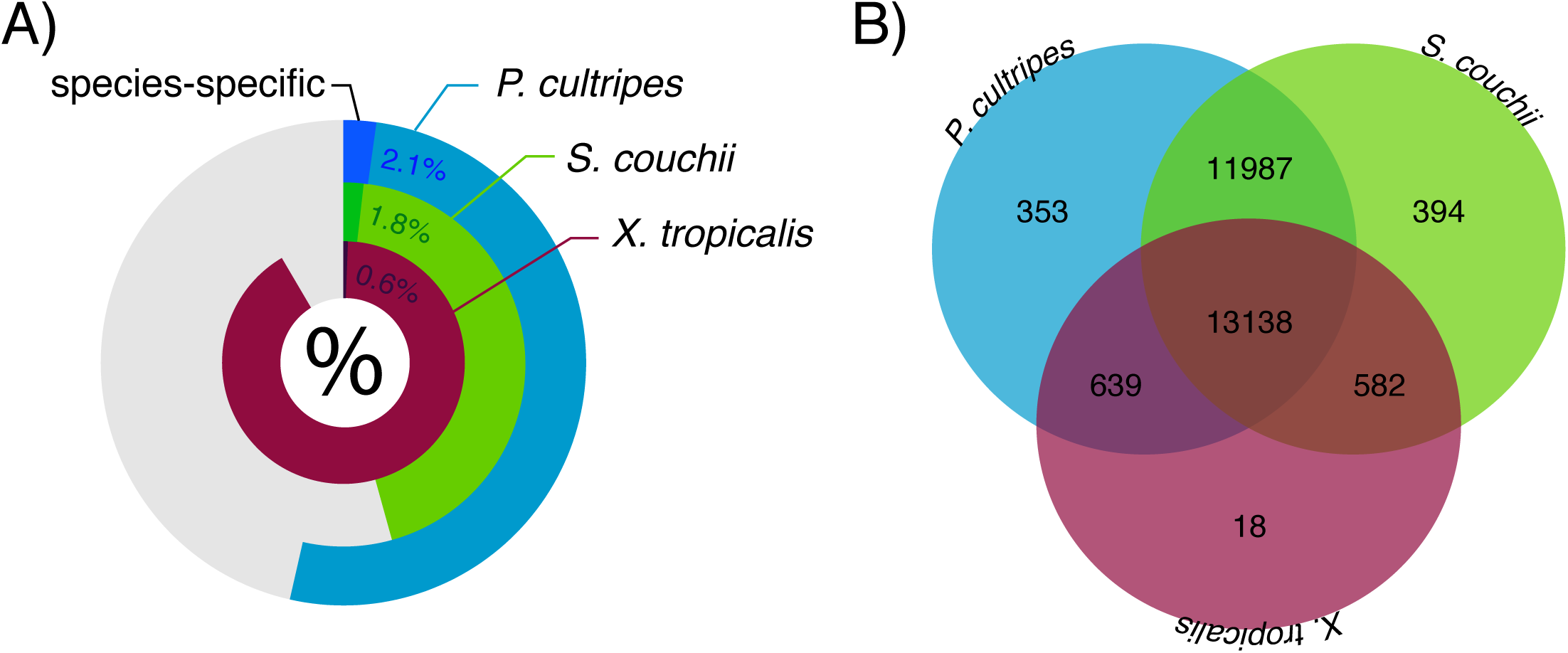
Orthofinder results showing a) the percentage of genes that could be assigned to orthogroups per species (darker shading represents percentage of genes in species-specific orthogroups) and b) the number of species-specific and shared orthogroups recovered.

Of the assigned genes, only small fractions of genes were in species-specific orthogroups (*X. tropicalis*: 0.6%, *P. cultripes*: 2.1%, *S. couchii* 1.8%; Figure 4a). Fifty percent of all genes were in orthogroups with two or more genes (G50 was 2) and were contained in the largest 23,404 orthogroups (O50 was 23 404). There were 13,138 orthogroups with all species present (Figure 4b) and 1,345 of these consisted entirely of single-copy genes. *Pelobates cultripes* and *S. couchii* shared substantially more orthogroups than either did with *X. tropicalis* and with 12,734 orthogroups being unique to these two species and therefore potentially important additions to the knowledge base of amphibian transcriptomics.

## Conclusion

*De novo* transcriptome assemblies of the larval phase of two amphibians with vastly differing environmental sensitivity in developmental rate are presented and annotated. Despite having drastically different sized genomes (with that of *Pelobates cultripes* being 2.6 times larger than that of *Scaphiopus couchii*; Liedtke et al. *in press*), the assemblies are of similar sizes (0.58Gbp vs. 0.64Gbp). The assemblies are of high quality, with ∼94% of raw reads mapping onto the transcriptomes, and both transcriptome assemblies consist of >86% full length BUSCO matches with only 2.9% of the assemblies having no corresponding match.

The PANTHER results suggest the two transcriptomes are largely comparable in their annotations and how they differ from *X. tropicalis*, but the overrepresentation test did not identify unexpected species-specific differences. For example, the analysis revealed that genes related to response to stimulus are under-represented in *S. couchii*, this is particularly true for the subcategory ‘response to abiotic stimulus’ (GO:0009628), which may reflect the fact that the development of *P. cultripes* is known to be more environmentally sensitive [3].

Approximately 40% of the assemblies were predicted to be protein coding sequences allowing for extensive annotation and here we provide information on SwissProt proteins (and their GO terms), protein family proteins (Pfam; and their GO terms), protein orthologous groups (eggnog), biological pathways (KEGG database), signal peptide cleave sites (SignalP) and transmembrane protein predictions (TMHMM). The number of predicted coding sequences (CDS=154,906 and 175,331) far exceeds that for published amphibian larvae transcriptomes (*R. catesbeiana*: 51,720 CDS (15% of assembly) [15], *M. fissipes* 51,506 CDS (46.8% of assembly) [14], *R. marina* 62, 365 CDS [13]) with substantial spadefoot toad-specific clusters of orthologous genes. The herein provided transcriptomes should therefore serve as an important resource for the advancement in the understanding of amphibian larval transcriptomics.

## Data and materials

The data sets supporting the results of this article are available in the associated repository GigaDB repository [<accession numbers to be released>]. Specifically, we provide Quality assessment results of both BUSCO and BowTie2, Transcriptome annotations including Trinotate summary tables, Panther annotations and transdecoder.pep sequence files. In addition, all raw reads as well as the transcriptome assemblies are deposited on the NCBI’s Sequence Read Archive [SRA; SRP161446] and Transcriptome Shotgun Assembly database [TSA; <accession numbers to be released>], under BioProject [PRJNA490256].

## Acknowledgements

This project was funded by Ministerio de Economía, Industria y Competitividad (MINECO) through the grant CGL2014-59206-P awarded to IGM. The CNAG team was funded by grant PT17/0009/0019 from MINECO, as well as Fondo Europeo de Desarrollo Regional (FEDER).

